# Arabidopsis ECERIFERUM3 (CER3) Plays a Critical Role in Maintaining Hydration for Pollen-Stigma Recognition during Fertilization

**DOI:** 10.1101/724559

**Authors:** Faqing Xu, Xiaojing Li, Zhongnan Yang, Sen Zhang

## Abstract

Plants distinguish the pollen grains that land on their stigmas, only allowing compatible pollen to fertilize female gametes. To analyze the underlying mechanism, conditional male-sterile mutations with affected pollen coat and disrupted pollen-stigma recognition were isolated and described. The mutant pollen failed to germinate, but germinated in vitro, suggesting that they are viable. In mutants, stigma cells that contacted their own pollen generated callose, a carbohydrate produced in response to foreign pollen. High humidity restored pollen hydration and successful fertilization, indicating defective dehydration in pollen-stigma interaction. Further analysis results from mixed pollination experiments demonstrated that the mutant pollen specifically lacked a functional pollen-stigma recognition system. The sterile plants lacked stem waxes and displayed postgenital fusion between aerial floral organs. In addition, the mutant pollen was deficient in long-chain lipids and had excess tryphine. Transmission electron microscopy observation showed that mutant pollen had almost the same surface structure as the wild type at bicellular pollen stage. However, abnormal plastoglobuli were observed in the plastids of the mutant tapetum, which was indicative of altered lipid accumulation. *CER3* transcript was found in anther tapetum and microspores at development stage 9 while CER3-GFP fusion protein was localized to the cell plasma membrane. Our data reveal that CER3 is required for biosynthesis of tryphine lipids which play a critical role in maintaining hydration for pollen-stigma recognition during fertilization.

## Introduction

In flowering plants, fertilization is a series of complicated processes involving many cellular interactions between pollen and stigma until the formation of zygotes following the fusion of male-female gametes. These cellular interactions determine interspecific or self-incompatibility during fertilization where the female reproductive tissues recognize and reject the pollen grains from closely related species or from the same individual plant depending on different plant species. Crucifer species are self-incompatible. Pollen development following self-pollination is arrested primarily before the pollen grains germinate. In compatible pollination, the stigma releases water and other substances to the mature pollen grain after it is deposited on the stigma. This allows the pollen grain to germinate. During germination, a pollen tube grows quickly through the transmitting tract of the style, delivering sperms to ovules within the pistil where fertilization takes place (Swanson et al. 2004). In incompatible pollination, the stigma does not release water responding to the landing of incompatible pollen grains which, therefore, are unable to germinate. Pollen hydration on dry stigma is an important and highly regulated step involved in blocking self-incompatible pollination (Hülskamp et al. 1995; Sarker et al. 1988) and in rejecting foreign pollen in interspecific crosses (Dickinson et al. 1995). While the underlying mechanisms remain unclear, it was shown that several stigma and pollen coat components may play a role, including aquaporins, lipids, and proteins. For example, several putative *Arabidopsis* aquaporins are highly expressed in the stigma (Swanson et al. 2005). The *Arabidopsis* pollen coat is involved in mediating the early contacts between the pollen grains and the stigma. Pollen coat originates from the tapetum layer which surrounds the developing microspores and is responsible for producing the exine precursors and pollen coat components. The pollen coat protects pollen grains from excess desiccation after anther dehiscence, contributes to pollen adhesion to the stigma, and, most importantly, facilitates pollen hydration (Edlund et al. 2004). The pollen coat protein, oleosin-domain protein GRP17, is required for the rapid initiation of pollen hydration on the stigma (Mayfield and Preuss 2000). In addition, some extracellular lipases (EXLs; Mayfield et al. 2001) were found in *Arabidopsis* pollen coat and are required for efficient pollen hydration. Mutation in *EXL4* leads to slower pollen hydration on the stigma and decreased competitiveness in pollination relative to wild type (Updegraff et al. 2009). Lipases catalyze acyl transfer reactions in extracellular environments (Upton and Buckley 1995). Analyses of mutants in lipid biosynthesis indicate a role of lipids as signaling substances in mediating water release at the stigma. For instance, the *cer* mutants fail to hydrate on the stigma because of decreased lipid content in their pollen coat—a defect that can be overcome by high humidity or the application of appropriate lipids to the stigma (Preuss et al. 1993; Hülskamp et al. 1995; Wolters-Arts et al. 1998; Fiebig et al. 2004;). On the other hand, any changes of long-chain fatty acids in stigma cuticle also affect pollen hydration. An organ fusion mutant, *fiddlehead* mutant exhibits abnormal lipid content in the cuticle of vegetative tissue (Pruitt et al. 2000). *fiddlehead* mutant lacks a β-ketoacyl-CoA synthase that is involved in the synthesis of long-chain fatty acids (Lolle et al. 1998). In this mutant, leaf cuticle permeability increases and pollen hydration is stimulated on inappropriate cell surfaces (Pruitt et al. 2000). Thus, the stigma cuticle may be adjusted for suitable water permeability in response to interaction with pollen grains (Pruitt et al. 2000).

In this study, we provide evidence that a conditional male-sterile *Arabidopsis* mutation in *CER3* alters the tryphine structure of the pollen surface. A T-DNA insertion disrupted the expression of *CER3* and resulted in excess lipids accumulation in the tapetum and pollen coat. The mutant pollen was viable but no longer communicated properly with the stigma; pollen germination failed as a result of limited pollen hydration and synthesized callose in stigma cell wall responding to the contact of mutant pollen. High humidity or co-pollination with the wild-type pollen led to successful fertilization of *cer3-8* mutant.

## Methods

### Plant materials

Seeds of *Arabidopsis thaliana* Col-0 were sown on the vermiculite and allowed to imbibe for 3 days at 4 ℃. Plants were grown in long-day conditions (16 h of light / 8 h of dark) in a growth room of about 22 ℃. The *cer3-8* and *cer3-9* mutant plants were isolated from the Col ecotype lines CSA306491 and CS306525 from the *Arabidopsis* Biological Resource Center (Columbus, OH), respectively. Before phenotype analysis, *cer3-8* and *cer3-9* plants were back-crossed to wild-type Col-0 three or four times.

### Phenotype characterization and microscopy

Plants were photographed with a Canon digital camera (Powershot-A710IS). Alexander staining was performed as described (Alexander 1969). Cross sections of anthers were performed as described (Zhang et al. 2007). Scanning electron microscope and transmission electron microscope of microspores and anthers were performed as described (Zhang et al. 2007). Photographs were taken with an Olympus BX51 microscope or a Carl Zeiss confocal laser scanning microscope (LSM 5 PASCAL).

For callose staining, emasculated wild type and *cer3-8* flowers were hand-pollinated with pollen grains from Col-0 wild type and *cer3-8* plants, respectively. Then the pistils were removed and placed on the slide and stained with aniline blue solution (0.1g / L in 50 mM K_3_PO_4_ buffer, pH 7.5). The stained pistils were covered with cover glass for observation under an Olympus BX51 fluorescence microscope.

For the pollen in vitro germination assay, mature pollen grains were spread onto medium consisting of 18% (m / v) sucrose, 0.01% (m / v) boric acid, 1 mM CaCl_2_, 1 mM Ca(NO_3_)_2_, 1 mM MgSO_4_, and 0.5% (m / v) agar (Li et al. 1999) in a humid chamber at approximately 22 °C. Single images were obtained with the Olympus BX51 microscope.

To analyze the hydration of pollen grains following hand-pollination, the pollinated flowers were allowed to develop for 20 minutes, the pistils were removed and placed on a slide, and the tissue were examined using an Olympus BX51 microscope. Hydration was assessed by the change from the elliptical shape of mature pollen grains to a nearly spherical shape. In the case of grains that failed to hydrate no change was observed.

### Analysis of fatty acids by GS/MS

For pollen coat extraction, *Arabidopsis* inflorescences were clipped, washed in phosphate buffer and filtered through cheesecloth. The filtered solution was centrifuged to obtain a pollen pellet. The pollen coat was then removed by washing the pollen grains three times in 10 volumes of cyclohexane based on the method by Doughty et al., 1993 with few modifications. Briefly, coating was removed from pollen grains by adding 800 µl of cyclohexane to 75 mg of pollen and agitating until suspended (5 sec). After separation by centrifugation (14,000 g, 20 sec), the cyclohexane fraction was transferred to a clean and dry eppendorf tube. Fatty acids methyl esterization was performed as previously described (Browse et al. 1986) with some modifications. Before analysis, 10 µl of N-methyl-N-trimethylsilyl-trifluoroacetamide (Fluka) were added and incubated at 37 ℃ for 30 min. As an internal control, 50 µl of nonadecanoic acid methyl ester (Fluka) stock solution (2 mg / ml in cyclohexane) was added. GC-MS was performed using Agilent 5975 inert GC / MS system with an HP-INNOWax column (Agilent).

### Treatment of pollen grains with formaldehyde

Wild-type flowers were removed from plants at the time of anthesis and placed on a glass slide. The slide was placed over a puddle of 37% aqueous solution of formaldehyde inside a culture dish for the stated time period. Anthers from these flowers were then used immediately in pollen rescue experiments.

### PCR and molecular cloning of the CER3 gene

The T-DNA insertion sites in the mutants were verified using PAC161 vector left-border primer PAC161-LBa1 (5’-TCCCCTGATTCTGTGGATAACCG-3’) and SALK line CS306491 genome-specific primers: LP306491 (5’-CCTCAAAACATTCCTCAGCAG-3’) and RP306491 (5’-TTAATGCGATGAGTCCTTTCG-3’); and SALK line CS306525 genome-specific primers: LP306525 (5’-GGACTCATCGCATTAATTGTGC-3’) and RP306525 (5’-CGAATCTTCTTTTGGAGTTCCC-3’). Co-segregation of the T-DNA insertion site and mutant phenotype was analyzed with the above LP and RP primers. For mutant plants, PCR with LB-A1-PAC161 and LP primers could amplify DNA fragments of ∼1000-bp and ∼700-bp, respectively. For wild-type plants, only PCR with LP and RP primers could amplify a DNA fragment of about 1154-bp (or ∼880-bp). For heterozygous mutant plants, PCR with both primer pairs showed positive results.

For complementation experiment, a 3434-bp *CER3* genomic fragment was amplified using KOD polymerase (Toyobo) and the gene-specific primers CP-F, 5’-<u>GGTACC</u>TACCCAATGTTAAATGAATGCGG-3’ and CP-R, 5’-<u>TCTAGA</u>ATTTGTGAGTGAAGAAACAGCAC-3’ (*Kpn*I and *Xba*I sites were underlined, respectively). The PCR product was cloned into the pMD18-T vector (Takara). After verification by sequencing, the fragment was subcloned into the binary vector pCAMBIA1300-GFP (CAMBIA; http://www.cambia.org) resulting in construct pCAMBIA1300-CER3-GFP, driven by its native promoter. This construct was introduced into homozygous mutant plants using the floral-dip method (Clough and Bent 1998) with *Agrobacterium tumefaciens* strain LBA4404. Transformants were selected on 1 / 2 MS medium containing 20 mg / L hygromycin and screened for fertile plants with homozygous background. For homozygous background verification, as LP / RP-amplified sequences are included in the complementation fragment, the following primers were used: LB-A1-PAC161 / RP primers to validate the existence of the T-DNA insertion in *CER3*; LP / RP primers to detect either the *CER3* genomic sequence or the transgenic complementation fragment; and genome-specific primers (CHZJD-F, 5’-TCTAGGCCTCTACTCGTCACAAT-3’; CHZJD-R, 5’-AGGAGATGAGTGGTGGAAAGAGT-3’) validate the homozygous background. As CHZJD-F was designed 445-bp upstream of CP-F, PCR with the CHZJD-F / CHZJD-R primer set was not able to amplify a 2.7-kb fragment in homozygous plants even if the complementation fragment was integrated into the genome.

### Subcellular localization of CER3

The above complementary construct pCAMBIA1300-CER3-GFP was also used for CER3 subcellular localization experiment. This construct was transiently expressed in tobacco leaf cells via *Agrobacterium tumefaciens* strain LBA4404 as described (Hu et al. 2002).

### Reverse transcription (RT)-PCR

Total RNA extraction, cDNA synthesis and RT-PCR analysis were performed as described (Zhang et al. 2007). The primers used for RT-PCR in analyzing expression level of *CER3* in *cer3-8* and *cer3-9* mutants were as follow: RT-F (5’-ATGGTTGCTTTTTTATCAGCTTG-3’) and RT-R (5’-ATTTGTGAGTGAAGAAACAGCAC-3’).

### in situ hybridization

Non-radioactive RNA *in situ* hybridization was performed using a digoxigenin (DIG) RNA labeling kit and PCR DIG probe synthesis kit (Roche; http://www.rochediagnostics.us) according to the manufacturer’s instructions. A 450-bp *CER3* cDNA fragment was amplified using *CER3*-specific primers: forward 5’-<u>GAATTC</u>TTGACATTGTCTATGGAAAG-3’ and reverse 5’-<u>GGTACC</u>ATTTGTGAGTGAAGAAACAG-3’ (*Eco*RI and *Kpn*I sites were underlined, respectively); The PCR product was cloned into the pBluescript SK vector (Stratagene; http://www.stratagene.com) and confirmed by sequencing. Plasmid DNA was completely digested by *Eco*RI or *Kpn*I, and used as a template for transcription with T3 or T7 RNA polymerase, respectively. Images were obtained with an Olympus DP70 digital camera.

## Results

### cer3 pollen cannot germinate on the stigma, but germinate in vitro

By screening of T-DNA tagged lines from the *Arabidopsis* Biological Resources Center (ABRC), two lines CS306491 and CS306525 were isolated which exhibited sterile phenotypes (Figure 1A). These mutants grew and developed normally as wild-type plants and pollen could release onto the stigma surface (Figure 1, B—D) while no seeds were produced. Both mutant plants produced seeds when fertilized with wild-type pollen, while application of mutant pollen to either wild-type or mutant plants produced no seeds. Hence, the mutation impaired only the reproductive system. Both mutant heterozygotes were fertile and one-fourth of their progeny showed a sterile phenotype. Thus, their fertility defects were caused by a recessive mutation in a single genetic locus, respectively.

**Figure 1.**
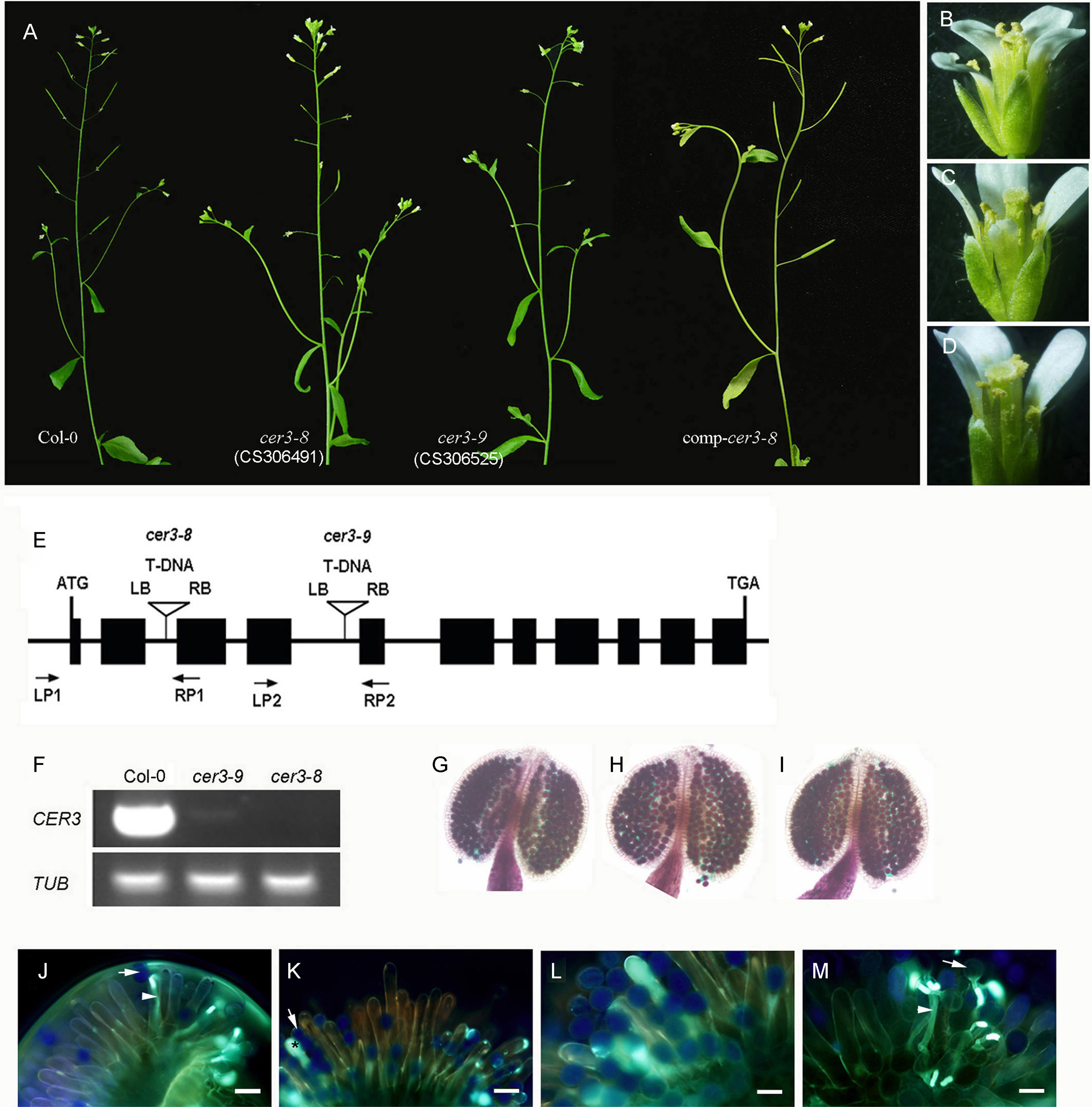
Phenotype characterization of *CER3* mutant alleles. (A) Wild type, *cer3-8*, *cer3-9* and *cer3-8* complemented plant images. (B) Wild-type (Col-0) flower. (C) *cer3-8* flower. (D) *cer3-9* flower. (E) Schematic representation of the genomic region of *CER3* gene. Boxes represent exons and lines indicate introns (not to scale). Triangles indicate the T-DNA insertion sites of *cer3-8* and *cer3-9* plants in the *CER3* gene, respectively. Arrows indicate oligonucleotide primer pairs LP1/RP1 and LP2/RP2 used for molecular characterization of the T-DNA loci. (F) Expression analysis of the *CER3* gene in wild-type (Col-0), *cer3-8* and *cer3-9* plants.*TUB*, *TUBULIN*. (G) A wild-type anther with viable pollen grains (stained). (H) A *cer3-8* anther with viable pollen grains (stained), (I) A *cer3-9* anther with viable pollen grains (stained). (J) Wild-type stigma pollinated with wild-type pollen and stained with analine blue, showing fluorescent pollen tube (arrowhead). (K) Wild-type stigma pollinated with *cer3-8* pollen and stained with analine blue, showing fluorescent callose (asterisk). (L) *cer3-8* stigma pollinated with *cer3-8* pollen and stained with analine blue, showing fluorescent callose (asterisk). (M) *cer3-8* stigma pollinated with wild-type pollen and stained with analine blue, showing fluorescent pollen tube (arrowhead). Arrows show pollen grains in j to m. Bars = 40 µm in (J and K); bars = 20 µm in (L and M).

To identify the corresponding loci responsible for the sterile phenotype, the genomic DNA fragments flanking the borders of T-DNA were recovered by PCR. T-DNA / plant genome DNA junctions can be amplified with the T-DNA left border primer, PAC161-LBa1, and genome specific primers. Sequencing of the PCR products showed that the T-DNAs were inserted at the 88th nucleotide in the second intron of the *CER3* gene in line CSA306491, and at the 248th nucleotide in the fourth intron of the *CER3* gene in line CS306525, respectively (Figure 1E). RT-PCR analysis showed that almost no transcript of *CER3* was observed in *cer3-8* mutant, while the expression level of *CER3* was greatly reduced in *cer3-9* mutant as compared to that in the wild-type (Figure 1F). Allelic test analysis indicated that both mutants were *CER3* mutation alleles. So lines CSA306491 and CS306525 were renamed as *cer3-8* and *cer3-9*, respectively. *cer3-8* mutant was used for further study here unless otherwise specified.

Genetic complementation experiment was performed with wild-type *CER3* genomic fragment fused in the modified pCAMBIA 1300 vector. Totally 10 transformants were obtained, and PCR analysis results showed that all the transgenic plants were homozygous *cer3-8*. These plants were all fertile with long siliques (Figure 1A). These data indicated that mutation of *CER3* is responsible for the sterile phenotype. To explore the mechanism of male sterility of *cer3* mutants, both mutant anthers were analyzed with Alexander staining method (Alexander, 1969). The results showed that the pollen of both mutant and wild type plants were stained purple, indicative of viable pollen (Figure 1, G─I). Further experiment was performed to examine the germination ability of the mutant pollen. In wild type plants, pollen at the stigma surface usually germinates a pollen tube to deliver sperms to the ovules. Along with the tube growth, certain amount of callose is produced (Figure 1J) (Eschrich and Currier 1964). However, no pollen tubes were observed on self-pollinated mutant stigmas (Figure 1L), neither when the mutant pollen was applied to wild-type stigmas (Figure 1K). Thus, the defect in early pollen germination may account for the observed male sterility in the mutant plants.

Interestingly, callose was found on the stigma surface with the phenotype associated with the *cer3-8* defect. Stigmatic papillae in direct contact with *cer3-8* pollen (Figure 1K), but not wild-type pollen (Figure 1J) were highly fluorescent when stained with aniline blue, indicating that the mutant pollen stimulated callose formation in the stigma cell. This phenotype was observed in all sterile segregants from *cer3-8*/*+* heterozygotes and, thus, was attributed to the *cer3-8* mutation. The abnormal callose formation responding to the contact of mutant pollen was also observed when the mutant pollen was applied to the wild-type stigma but not when wild-type pollen was applied to mutant stigma (Figure 1, K─M). Thus, mutation of *CER3* makes pollen fail to germinate on the stigma and induces callose formation within the stigma cells.

Pollen from most plant species germinates effectively when cultivated in a medium containing sucrose, calcium, magnesium and borate (Li et al. 1999). Interestingly, when *cer3-8* pollen was cultivated in this medium, the pollen germinated almost the same as wild type (Figure 2). These results indicated that *cer3-8* pollen is viable and can produce, in vitro, all of the substances needed for pollen germination and tube growth.

**Figure 2.**
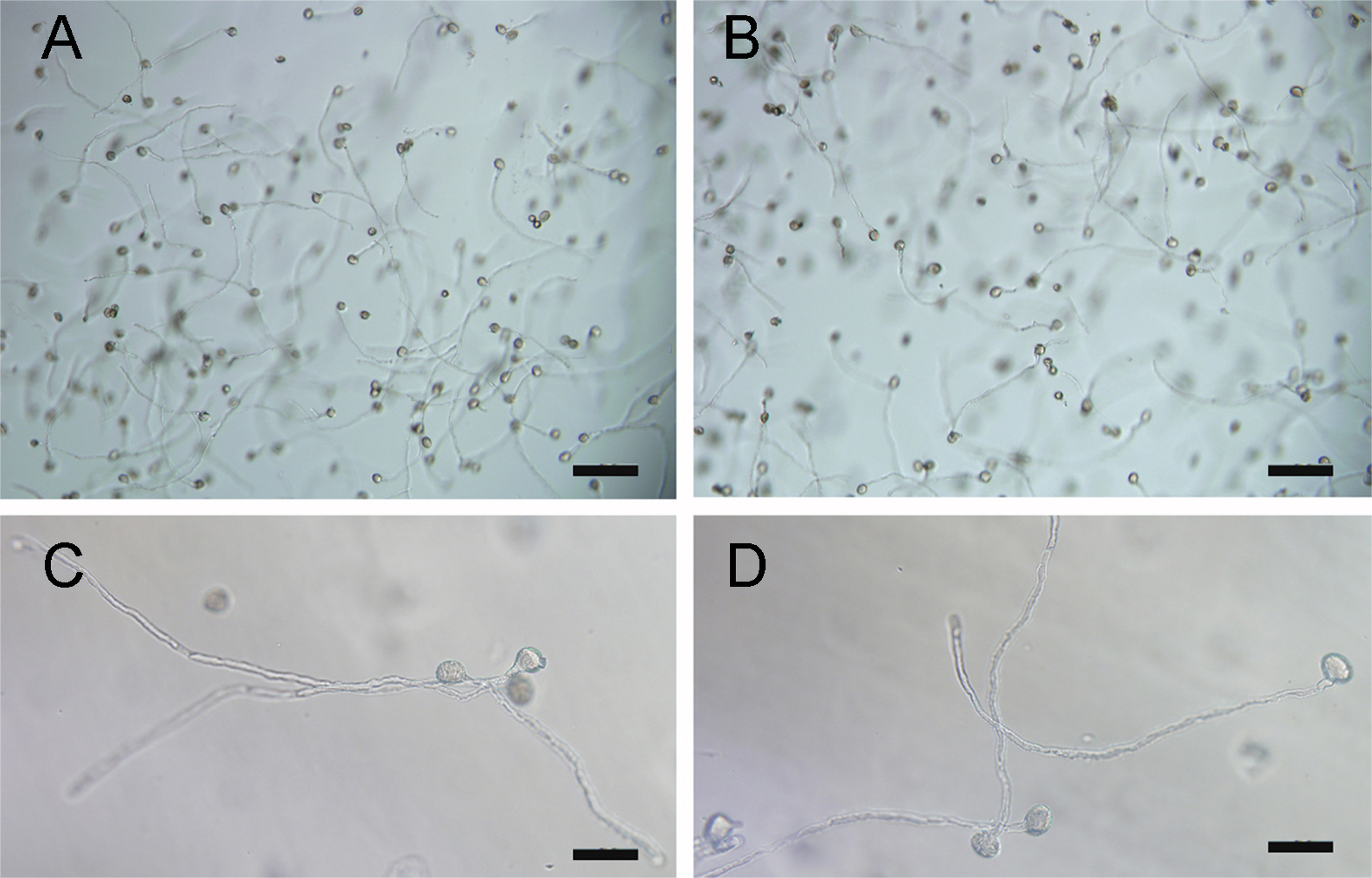
Germination of wild-type and *cer3-8* pollen in vitro. Pollen tubes were observed after wild-type (A) or *cer3-8* (B) pollen grains were incubated in pollen germination medium. (C and D) close-up views of (A and B), respectively. Bars = 100 μm in (A and B); bars = 1500 μm in (C and D).

### cer3-8 mutant pollen cannot hydrate on the stigma

To explore the nature of pollen not germination on the stigma of the *cer3-8* mutant, pollen hydration was first checked on the hand-pollinated stigmas. Wild-type stigmas were hand-pollinated and a few minutes later the stigmas were examined. The pollen can be found clearly undergone hydration turninging into a spherical shape from the unhydrated (Figure 3, A─C). When the mutant pollen was placed on their own stigma, no hydration took place for longer period of time (Figure 3, D─F).

**Figure 3.**
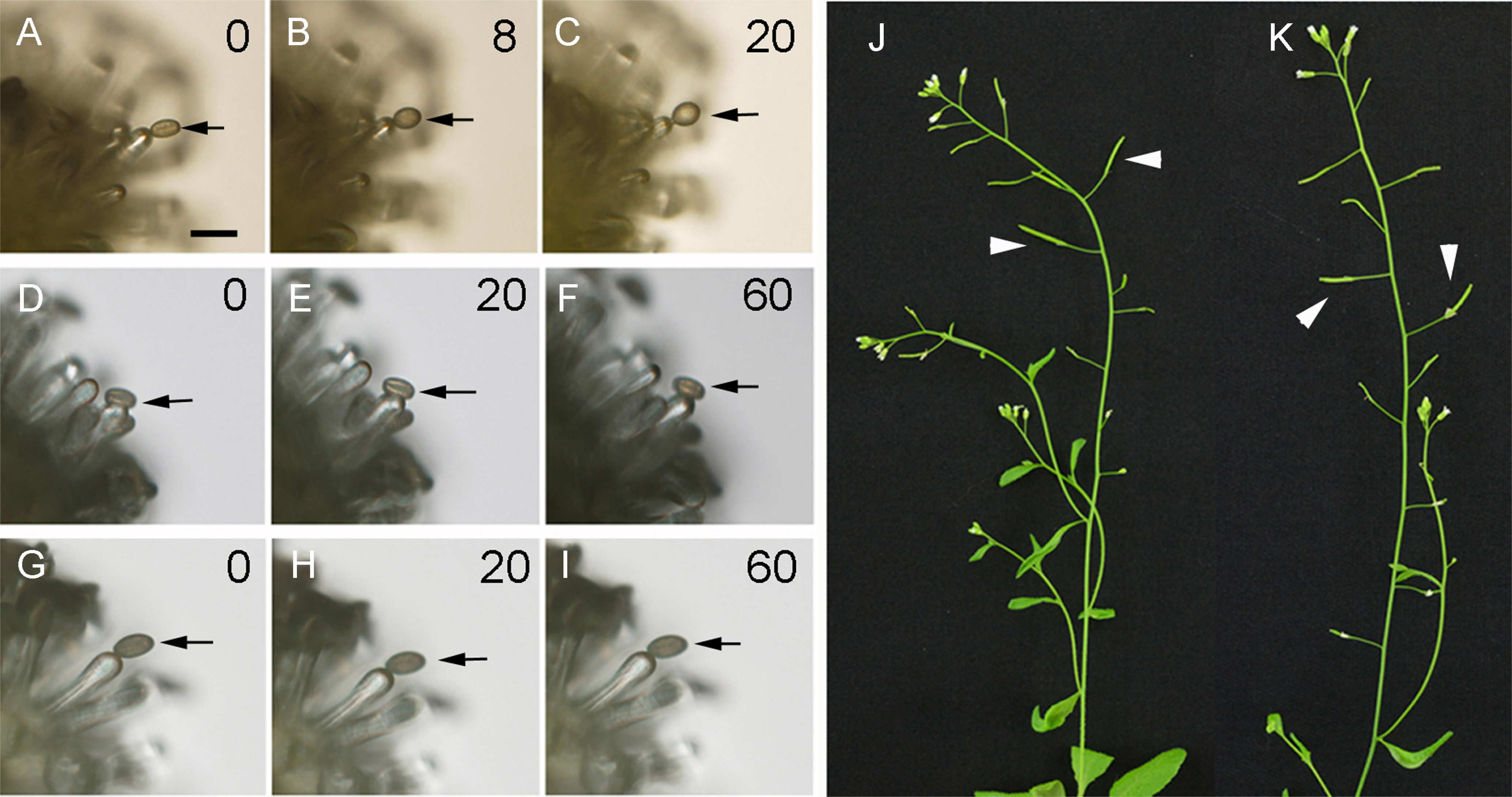
Hydration of wild-type, but not *cer3-8*, pollen occurred rapidly on the stigma surface (A-I), and high humidity resumed male fertility of *cer3-8* (J) and *cer3-9* (K). Wild-type pollen (A-C) expands within minutes when placed on a wild-type stigma surface. No change in pollen shape or size was observed in similar experiments with *cer3-8* pollen on a *cer3-8* (D-F) or a wild-type stigma surface (G-I) for longer period of time. Numbers (in the upper right corner) indicate time in minutes. Arrows indicate pollen grains. The examples depicted here are representative of similar observations of >100 pollen grains. Bars = 40 µm in (A-I). (J, K) Humidity restored fertility in *cer3-8* and cer3-9 plants, respectively. The plants grown in 50-70% relative humidity transferred to 90% relative humidity. Expanded seed pods (indicated by arrowheads) show that fertilization has occurred.

Further examination indicated that no hydration took place even when the mutant pollen was put on the wild-type stigma (Figure 3, G─I). Additional assays were performed with mutant pollen to determine whether they were capable of hydration after longer periods on the stigma, however no hydration was observed even several hours after pollination of 208 pollen, while 189 wild-type pollen were all hydrated.

The *cer3-8* pollen not hydration on the stigma, coupled with the germination of the mutant pollen in vitro, suggested that the *cer3-8* fertilization defect might be overcome by artificially wetting the pollen. To test this hypothesis, *cer3-8* plants were moved from normal growth condition (50-70% relative humidity) to a high-humidity environment (90% relative humidity). Fertility was restored as indicated by the expanded siliques in the mutant plants (Figure 3, J and K). The hydration of pollen was presumably facilitated by the passive absorption of water vapor from the environment (Figure S1). Thus, the effects on pollen function caused by *cer3-8* mutation are considered conditional and reversible.

### cer3-8 mutant specifically lacks recognition competence

The experiments described above suggested that the stigma can recognize different pollen, allowing hydration of wild-type but not *cer3-8* pollen. These data implied that the mutation affected the signal that was carried by the pollen and was required for the recognition of the pollen by the stigma. Thus, to further analyze the requirements for pollen hydration, *cer3-8* flowers were co-pollinated with wild-type and *cer3-8* pollen, by carefully placing the pollen side by side. As expected, viable seeds were produced. Interestingly, these seeds not only yielded fertile, *cer3* / + plants, but also infertile, *cer3* / *cer3* plants, indicating that the wild-type pollen elicited *cer3-8* pollen hydration. Pollination with a mixture of the two types of pollen (1 : 1) results in 39.5% homozygous mutant plants and 60.5% heterozygotes, suggesting that the rescue effect of mutant pollen was fairly effective. These results indicated that interaction between the wild-type pollen and the stigma can result in hydration of nearby mutant pollen.

In the above pollen rescue experiment, the rescuing pollen not only hydrate, but also germinate and enter the stigma surface. In order to determine the process involved in the rescue experiment, further analysis was carried out. Wild-type pollen was made inviable by treatment with formaldehyde vapor and then mixed with *cer3-8* pollen. Wild-type pollen treated in this way can hydrated on the stigma but did not germinate and develop further. The mixed pollen was placed on the *cer3-8* stigma, and the number of seeds produced per flower was determined (Table 1). The results indicated that the mutant can be rescued by the wild-type pollen that were capable of eliciting water transfer but were unable to germinate and enter the stigma surface. These results clearly demonstrate that the defect in the mutant is only limited to the hydration step of fertilization.

**Table 1.**
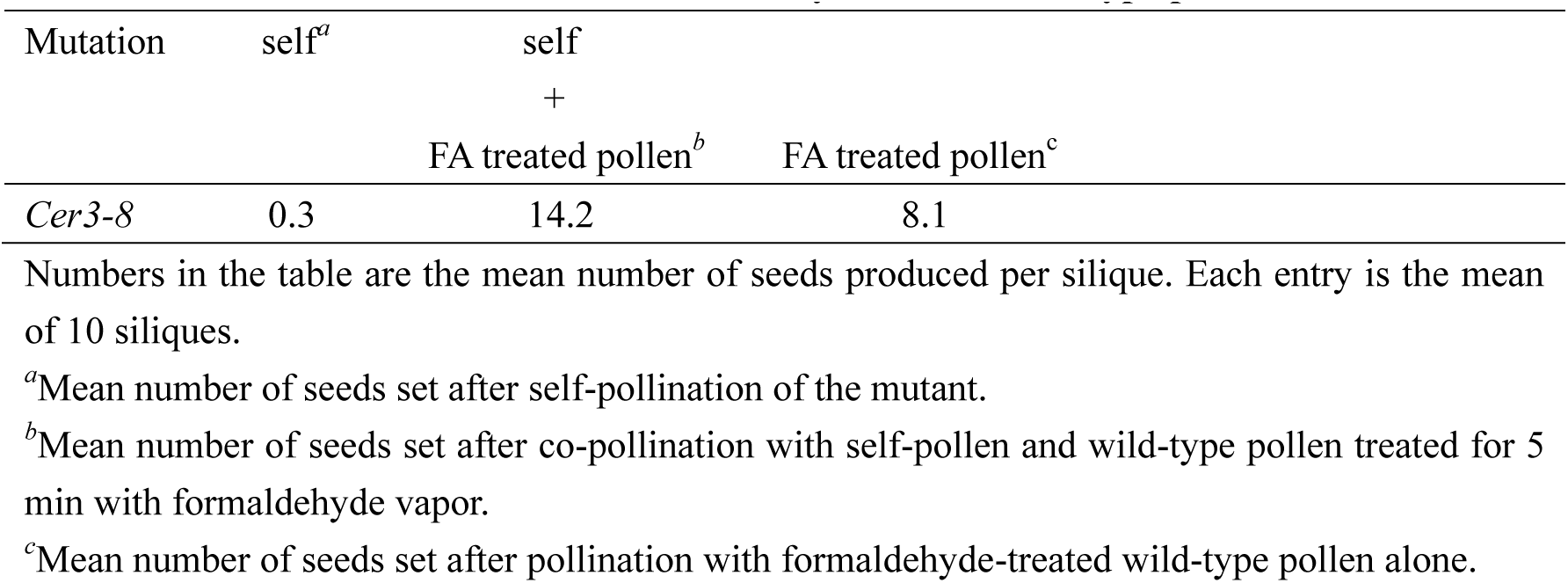
Rescue of cer3-8 mutant with formaldehyde treated wild-type pollen

### cer3-8 mutant is deficient in wax production with organ fusion

In addition to its male sterility, the *cer3-8* mutant was also defective in the production of waxes on the stem surface. Wax is composed of long-chain lipids and is visualized easily on the surface of wild-type stems as a dull, glaucous covering. By contrast, stems from the *cer3-8* mutant looked bright green and glossy in appearance (Figure 4A), resembling other wax-defective, *cer* mutants (Dellaert et al. 1979; Koomneef et al. 1989). Besides, postgenital fusions were observed between some aerial organs in the *cer3-8* mutant (Figure 4). Specifically, fusions were found to occur among stamens and styles and sepals (Figure 4B), and between different flower petals (Figure 4C). These results suggest that mutation of *CER3* affects the production of wax on the stem surface and some floral organs.

**Figure 4.**
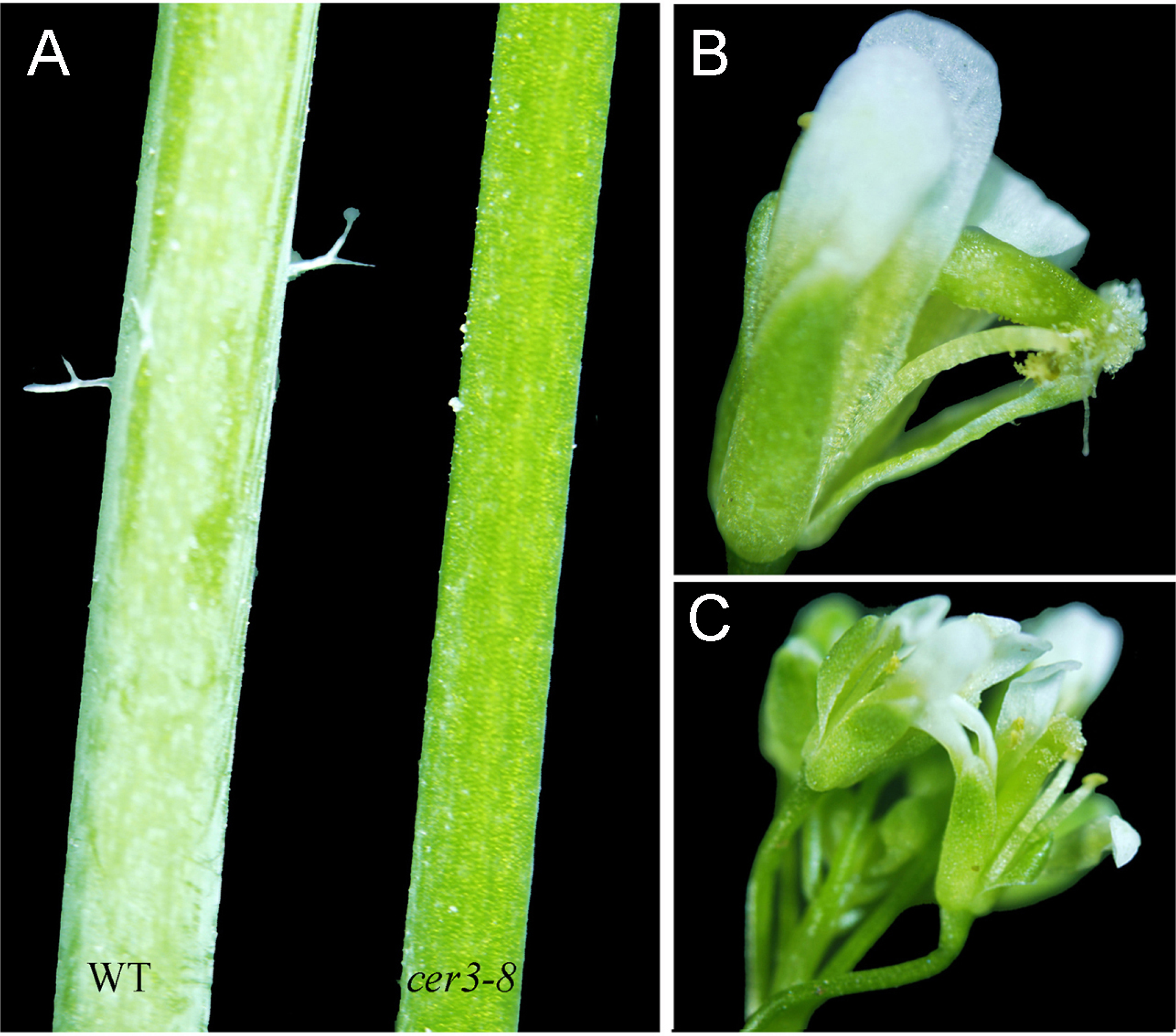
c*e*r3*-8* mutants are defective in epicuticular wax production, and postgenital fusion occurs between some floral organs. (A) *cer3-8* plants are deficient in the waxes that coat wild-type stems. Postgenital fusion does not occur in wild type flowers (Figure 1 B). Postgenital fusions occur between the stamen and style and sepal (B) as well as between petals (C) of different flowers in the mutants.

### cer3-8 pollen is deficient in long-chain lipids

As described above, the *cer3-8* mutants are defective in wax production and pollen germination, suggesting that lipids might play a role in pollen-stigma signaling. The lipid content of wild-type and *cer3-8* pollen coat was characterized and compared to examine whether long-chain lipids were present in the *cer3-8* mutant. Pollen coat lysates were prepared and extracted with cyclohexane, and the components in the organic phase were subsequently separated and analyzed by gas chromatography-mass spectrometry (GC-MS). All lipids detected by GC-MS were compared between mutant and wild type extracts (Figure 5), and several long-chain lipid compounds in wild-type pollen coat were missing or accumulated less in the mutant. The identity of these molecules was confirmed by analysis of their mass spectra, and their relative abundance is shown in Table 2. Twenty-nine-carbon (C29) molecules (nonacosene, n-nonacosane) were easily detected in the wild-type sample, but only a small fraction of these lipids (0-1% of wild-type levels) was found in the *cer3-8* extract, and 30-carbon molecules were not present at all (Table 2). Although long-chain lipid molecules were low in abundance, there were more abundant medium-chain lipids (16 and 18 carbons) in the mutant pollen coat (Figure 5). These results indicated that *cer3-8* mutants cannot extend lipid chains to a length of 29-carbon atoms or longer.

**Figure 5.**
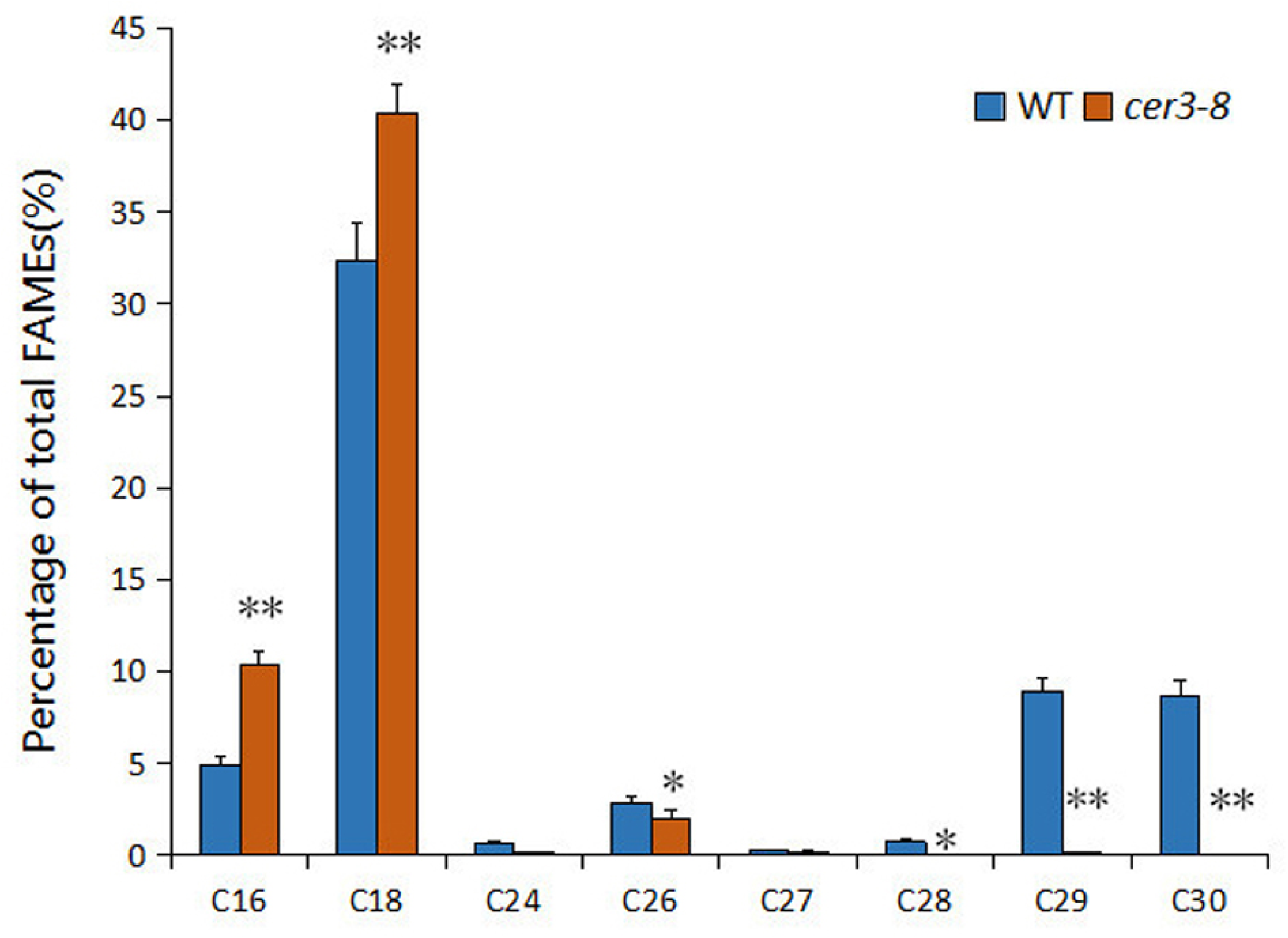
c*e*r3*-8* pollen is deficient in long-chain lipids. Pollen coat lysates were extracted with cyclohexane and analyzed by gas chromatography-mass spectroscopy. The wild-type extract contains 29- and 30-carbon lipids (identified in Table 2), whereas these compounds are virtually absent from the *cer3-8* extract. The data were analyzed from three biological replicates and presented as average SD. The statistics analysis was performed using student’s t-test (**p<0.01; *P<0.05).

**Table 2.** Lipid content in tryphine from cer3-8 and wild-type pollen

| Number of carbon atoms | Compound | Wild type | *cer3-8* |  |
| --- | --- | --- | --- | --- |
| 16 | hexadecanoic acid | 4.80 | 10.01 | (2.10) |
| 18 | octadecanoic acid | 31.77 | 39.0 | (1.23) |
| 26 | hexacosane | 2.78 | 1.91 | (0.69) |
| 29 | nonacosene | 5.91 | 0.06 | (0.01) |
| 29 | n-nonacosane | 2.83 | 0.0 | (--) |
| 30 | n-triacontane | 8.52 | 0.0 | (--) |

### cer3-8 mutant fertility can be restored by long-chain lipids

The above phenotype analyses of *cer3-8* mutation regarding pollen hydration defect and long-chain lipid deficiency in the pollen coat indicate the important role of long-chain lipids in pollen–stigma recognition during pollination. Thus, exogenous long-chain lipid melissic acid dissolved in chloroform was applied to a 37-day-old *cer3-8* mutant stigma surface (Figure 6C) with chloroform as control (Figure 6A). Elongated siliques were observed on the mutant plant treated with melissic acid 7 days later (Figure 6D), while siliques on the control plant were not changed (Figure 6B). These results demonstrated that fertility was restored by the exogenous application of long-chain lipids, suggesting that long-chain lipids are involved in pollen-stigma communication.

**Figure 6.**
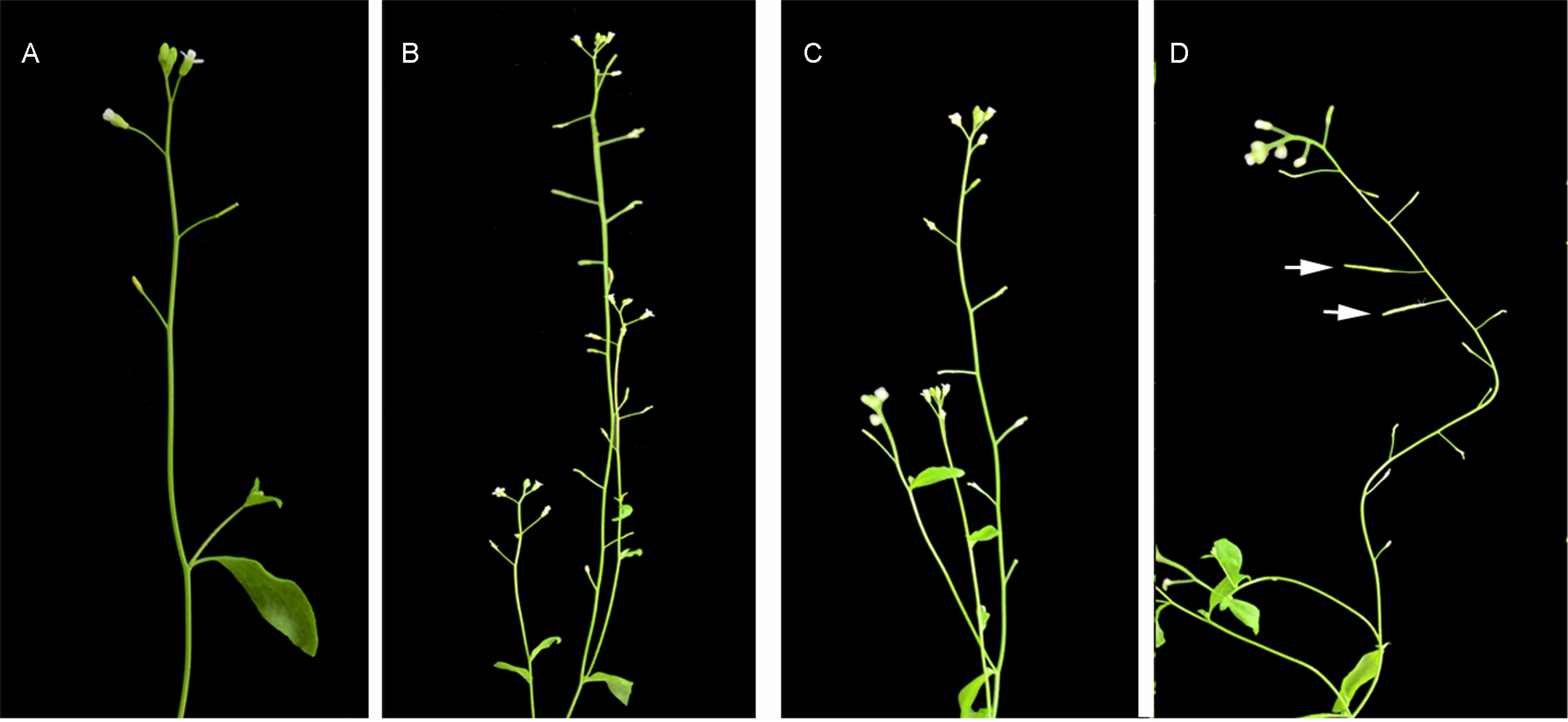
Fertility restoration of *cer3-8* Fertility of *cer3-8* was resumed as indicated by expanded siliques (showed by arrows in D) by application of melissic acid on the mutant floral buds (C) with application of chloroform on the mutant buds as control (A, B). Melissic acid dissolved in chloroform (25 µg / µl).

### cer3-8 mutation leads to smooth pollen surface

As described above, the effect of *cer3-8* mutation on male sterility is sporophytic, so this gene must be active in the diploid tissues. Its activity could be involved either in the pollen mother cell prior to meiosis or the tapetal cell surrounding the microspore. In order to study the surface structure of the *cer3-8* mutant pollen, the pollen was examined by scanning electron microscopy (SEM). Compared with wild type, many *cer3-8* pollen were abnormally stuck together (Figure 7, A and B), and the exquisite reticulate pattern of wild type pollen was not prominent in the mutant; instead excess coating was observed on the *cer3-8* pollen surface (Figure 7, C and D). SEM observation of FAA treated pollen further confirmed that some *cer3-8* pollen were stuck together (Figure 7, E and F), and surface of the *cer3-8* pollen was different from that of the wild-type (Figure 7, G and H).

**Figure 7.**
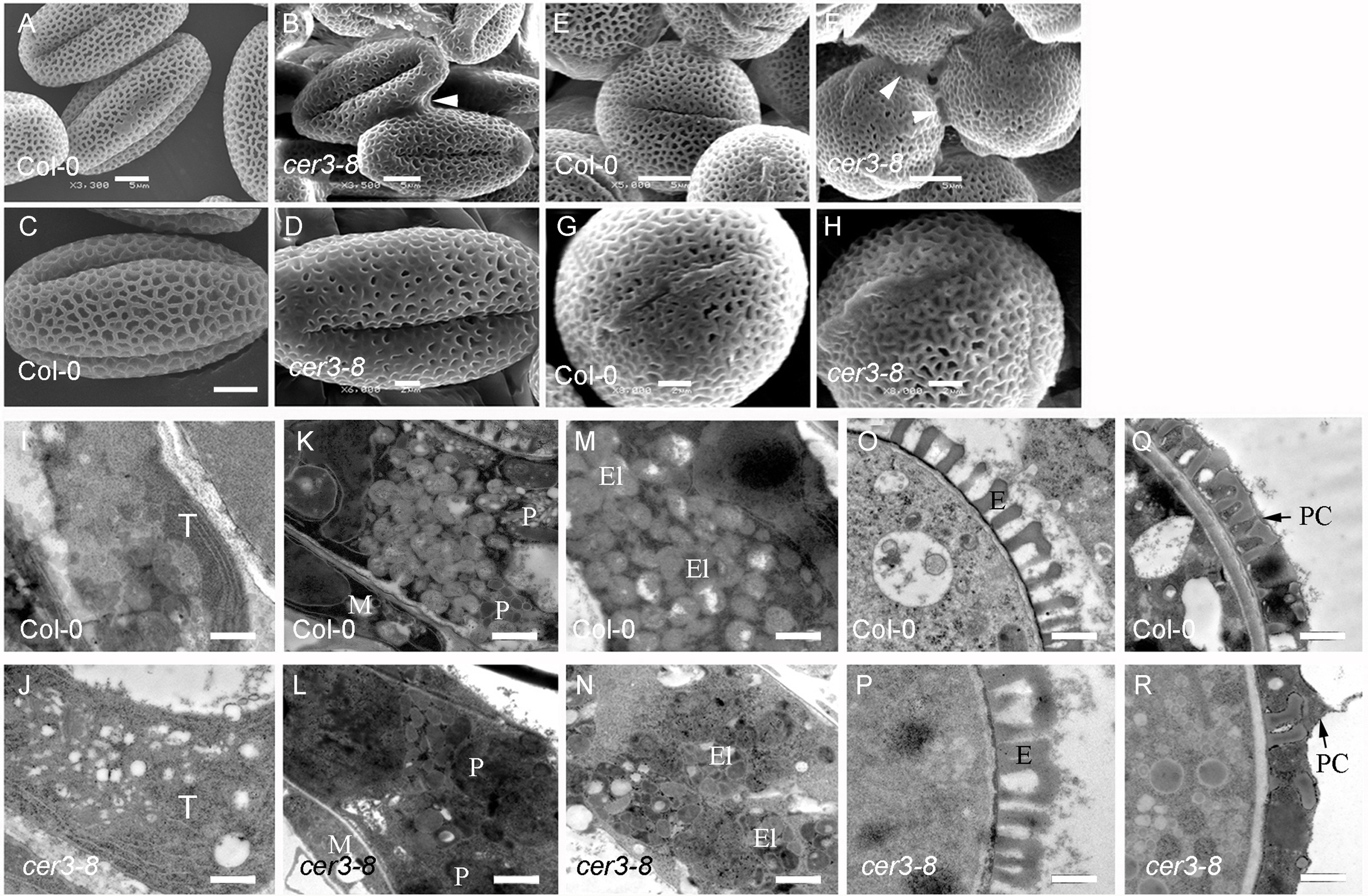
TEM analysis of the pollen surface (A-H), tapetum and pollen development in later stage anthers (I-R). (A) Wild-type pollen. (B) Some *cer3-8* pollen stuck together (arrowhead). (C) Wild-type pollen with exquisite reticulate pattern. Bar = 3 µm. (D) *cer3-8* pollen showing excess coating. (E) FAA treated wild-type pollen. (F) FAA treated *cer3-8* pollen stuck together (arrowheads). (G) Close-up view of wild-type pollen. (H) Close-up view of *cer3-8* pollen showing different surface from wild type. (I) Tetrad stage wild-type tapetum. (J) Tetrad stage *cer3-8* tapetum. (K) Uninucleate microspore stage wild-type tapetum, showing normal plastoglobuli. (L) Uninucleate microspore stage *cer3-8* tapetum, showing electron-dense plastoglobuli. (M) Bicellular pollen stage wild-type tapetum, showing translucent elaioplasts with compacted plastoglobuli. (N) Bicellular pollen stage *cer3-8* tapetum, showing less translucent elaioplasts with electron-densed plastoglobuli. (O) Bicellular pollen stage wild-type pollen. (P) Bicellular pollen stage *cer3-8* pollen showing similar exine to wild type. (Q) Tricellular pollen stage wild-type pollen. (R) Tricellular pollen stage *cer3-8* pollen with excess coating. Bars = 1 µm in (I to R). T, tapetum; M, middle layer; P, plastid; El, elaioplast; E, exine; PC, pollen coat.

In order to examine the origin of the excess pollen coating, transmission electron microscopy (TEM) experiment was carried out with developing anthers from the *cer3-8* and wild type plants. At the tetrad stage, numerous small and large electron-translucent vesicles were observed throughout the tapetum in the wild type and *cer3-8* plants (Figure 7, I and J). No obvious structural differences in the tapetum were observed at this time. At the uninucleate microspore stage in the wild type, the plastids contained numerous large, electron-transparent vesicles known as plastoglobuli (Figure 7K). In *cer3-8* mutant, the appearance of the plastids was similar to that in the wild type (Figure 7L). However, electron-dense granules were observed in the plastids of *cer3-8* mutant (Figure 7I; big arrow). At the bicellular pollen stage in the wild type, the plastids developed to relatively translucent elaioplasts consisting of compacted plastoglobuli (Figure 7M). However, the elaioplasts of *cer3-8* appeared to be less translucent, full of electron-dense granules indicative of lipid accumulation in the plastoglobuli, suggesting that the mutant plastids did not fully develop into elaioplasts (Figure 7N). At the same time no obvious differences were observed between the pollen exine of them (Figure 7, O and P). At the tricellular pollen stage in the wild type and the *cer3-8*, the tapetum was completely degenerated, and all the cell remnants were released and deposited on the maturing pollen surface. The *cer3-8* pollen was found to be covered with excess coating compared with that of the wild-type (Figure 7, Q and R). These data suggested that disruption of *CER3* may disturb the synthesis of precursors of tryphine from the tapetum to the developing pollen till the complete degeneration of the tapetum.

### CER3 is highly expressed in the tapetum and microspore

Previous studies have shown that *CER3* was expressed in siliques, stems, rosette leaves, cauline leaves, flower buds and open flowers, but not in roots (Ariizumi et al. 2003). To further study the functions of *CER3* during microspore development, RNA *in situ* hybridization experiment was performed to examine the precise spatial and temporal expression pattern of *CER3* during wild type anther development. The results showed that *CER3* transcript was initially detected at anther stage 7 and 8 tapetum (Figure 8, A and B). The highest hybridization signal was observed at stage 9 tapetum and microspores (Figure 8C). Then the signal was gradually reduced in the tapetum and microspores from stage 10 to 11 (Figure 8, D and E). By contrast, the control was barely detected at the stage 9 anther (Figure 8F). These results suggest that *CER3* is required for postmeiosis pollen development.

**Figure 8.**
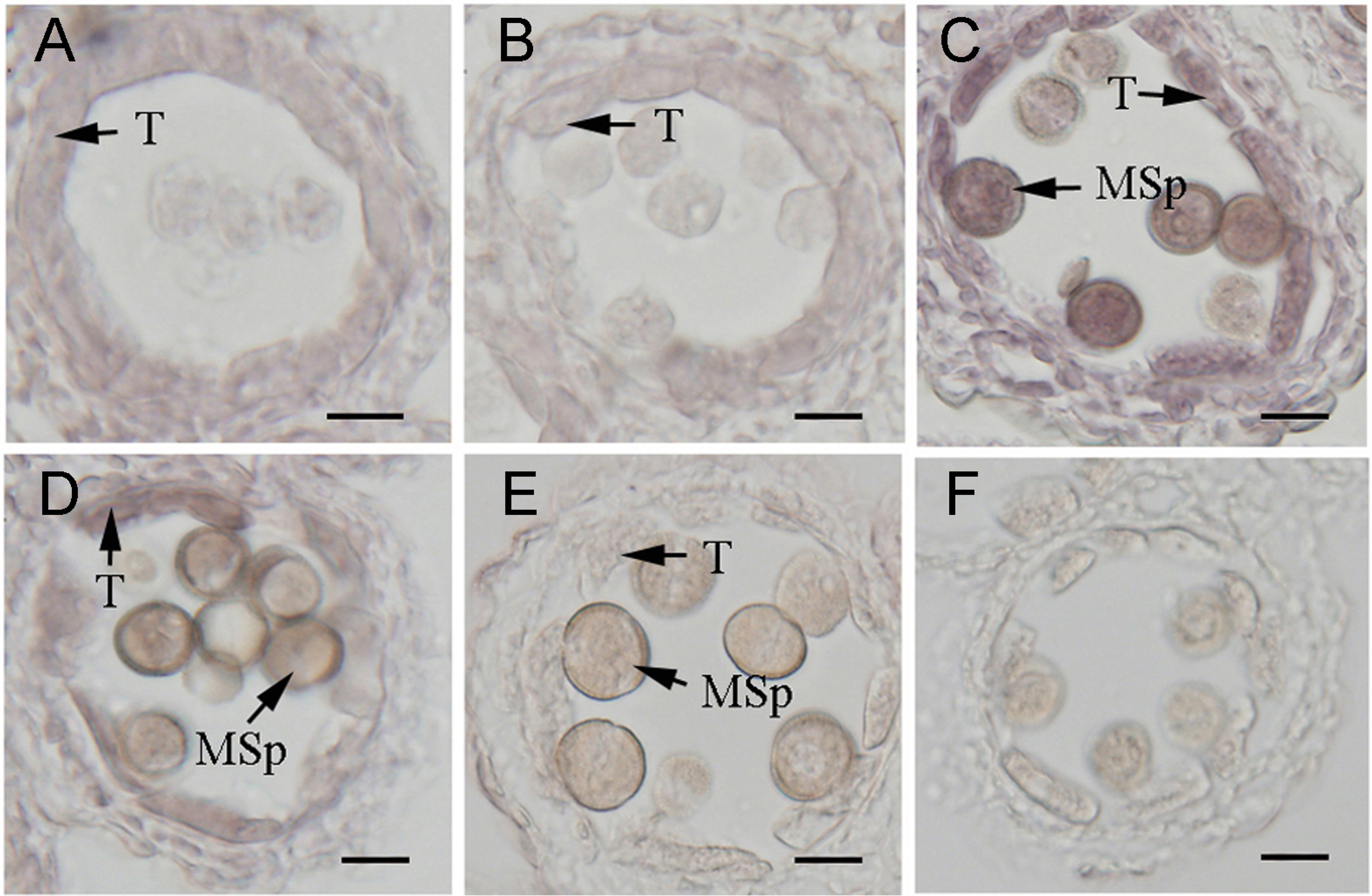
Expression pattern of *CER3*. Cross-sections through wild-type anthers at different stages of development probed with digoxigenin-labeled *CER3* antisense or sense probes. (A) Stage 7 anther, showing that *CER3* was slightly expressed in the tapetum. (B) Stage 8 anther, showing that *CER3* was slightly expressed in the tapetum. (C) Stage 9 anther, showing that *CER3* was strongly expressed in the tapetum and microspores. (D) Stage 10 anther, showing that *CER3* was highly expressed in the tapetum and microspores. (E) Stage 11 anther, showing that *CER3* was slightly expressed in the tapetum and microspores. (F) Sense probe showing almost no hybridization signal. Bars = 15 µm. T, tapetum; MSp, microspore.

### CER3 is a plasma membrane-localized protein

The former study has reported that the *CER3* gene encodes a protein of 632 amino acid residues which was predicted to be localized to the plasma membrane (Ariizumi et al. 2003). Our bioinformatics analysis confirmed six putative transmembrane regions in CER3 (Figure 9A) (Chen et al. 2003; Kurata et al. 2003). Experimentally, a *CER3*-GFP fusion driven by the *CER3* native promoter was generated in the modified pCAMBIA 1300 vector, and 35S-GFP was used as a control. These constructs were introduced into tobacco leaves as described (Hu et al. 2002). As shown in Figure 9, GFP was distributed throughout the cell expressing the control construct (Figure 9B), while the CER3-GFP was observed on the plasma membrane of the cell, indicating that CER3 is a plasma membrane-localized protein (Figure 9C).

**Figure 9.**
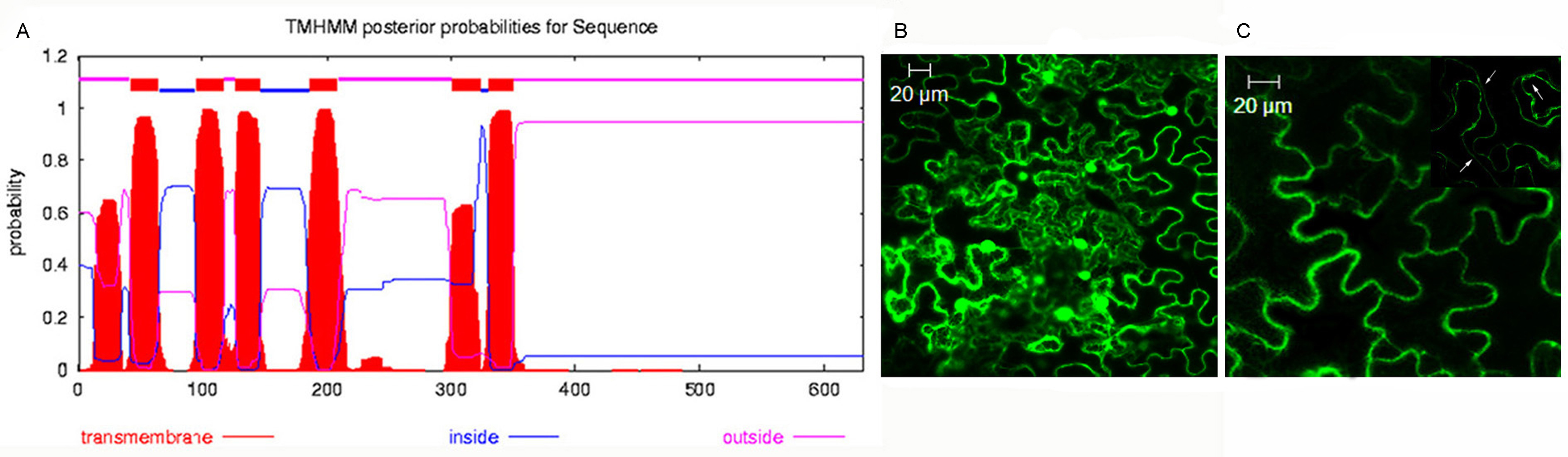
Subcellular localization of CER3. The transient expression in the infiltrated tobacco leaf cells was carried out for subcellular localization. (A) The transmembrane domain in CER3 protein was predicted using TMHMM (http://www.cbs.dtu.dk/services/TMHMM/). (B) Subcellular localization of 35S-GFP fluorescence in transient transgenic tobacco leaf cells. Green fluorescence was dispersed throughout the cell. (C) Subcellular localization of CER3:GFP fluorescence in transient transgenic tobacco leaf cells. Green fluorescence indicates the localization of CER3:GFP protein in the epidermis cell membrane. The insert shows plasmolyzed epidermis cells treated with 0.8 M mannitol, where arrows indicate plasmolysis.

## Discussion

The *cer3-8* mutants produce pollen that cannot send appropriate signals to the *Arabidopsis* stigma. At normal growth conditions, the mutant pollen does not hydrate and germinate on the stigma. Moreover, the appearance of callose in the stigma contacting the *cer3-8* pollen indicated an aberrant pollen-stigma interaction. In vitro germination data indicate that *cer3-8* pollen is viable. Besides, the defective fertility is recovered when the mutants grow in a high humid environment or when mutant pollen is co-pollinated with wild-type pollen. Thus, *cer3-8* represents a conditional, male-sterile mutation that specifically affects pollen function. Wax deficiency on the *cer3-8* plants indicates that long-chain lipids might be required for pollen function. Actually, the *cer3-8* mutant pollen lack long-chain lipids compared with the wild-type (Figure 5) in spite that *cer3-8* pollen were covered with excess coating. The excess covering was due to the final dumping release and deposition of the remnants abnormally accumulated from degenerated tapetum because of the disruption of *CER3* gene.-

The stigma papillae cells are receptive to pollen binding. Stigmas are usually classified into two types (wet and dry stigmas) based on the extracellular matrix that covers their surface. Wet stigmas are covered with sticky secretions, while dry stigmas are coated with a protein-containing pellicle (Heslop-Harrison and Shivanna 1977; Heslop-Harrison 1981; Heslop-Harrison 1992). The carbohydrate and lipid-rich matrix on the surface of wet stigmas may promote the hydration of most pollen. By contrast, dry stigmas may selectively promote pollen hydration (Roberts et al. 1980; Sarker et al. 1988; Preuss et al. 1993). Pollen hydration is the first step blocked in a self-incompatible pollination (Dickinson and Elleman 1985; Dickinson 1995). The analysis of wax-defective, *cer* mutations has demonstrated that the pollen coat is required for pollen hydration (Preuss et al. 1993; Hülskamp et al. 1995). The *CER* genes are known to function in wax biosynthesis (Hannoufa et al. 1993), producing long-chain lipids that cover the surface of the stems and leaves, as well as the pollen (Preuss et al. 1993). Mutations in these genes affect the amount of lipids in the pollen coat, and this deficiency may induce the loss of coat proteins and other components during development. Consequently, *cer* defects may influence an array of molecular interactions that normally occur between the pollen coat and the stigma.

So, the pollen coat lipid and protein are considered to be essential for pollen hydration. In contacting with wild-type pollen, the pollen coat could be changed, forming a contact zone between the stigma and the pollen (Elleman and Dickinson 1996). In this process, long-chain lipids and some other substances in the pollen coat are considered to signal and to be reorganized by the stigma through the actions of the lipid-binding proteins, and to create a capillary system through which water can flow from the stigma to the pollen (Murphy 2006).

The results of reduced or altered lipid composition on the pollen surface can be analyzed at the ultrastructural level. Previous study showed that a mutation in the *CER6* resulted in pollen without the tryphine layer (Preuss et al. 1993). Another partially sterile *CER6* mutant exhibited a reduced tryphine layer with reduced number and size of lipid drops. These results indicated that lipid products of the *cer* pathway are required as a binding agent to hold the tryphine layer to the pollen. In the tryphine layer lipid drops are missing in *cer1-147*, or are reduced in size in *cer6-2654* even though their pollen showed normal coat thickness (Hülskamp et al. 1995). These data implied that long-chain lipids may play an indirect role in solubilizing some other recognition factors present in the tryphine layer. Under the growth conditions used here, the *cer3-8* mutant produced stem and flower with less wax, pollen with excess tryphine that fails to germinate on the stigma surface, resulting in male sterility. The phenotype of *cer3-8* pollen coat structure was apparently different from the above described *cer* mutants. All of these results imply that lipids play important roles in the tryphine regarding pollen-stigma signaling and pollen hydration.

In wild-type *Arabidopsis* anther, the tapetum surrounds the microspore and provides materials for microspore development (Scott et al. 1991). At later stages in pollen development, the tapetum accumulates lipidic components in the pollen coat, which may be transported to the exine by transporters located at the tapetal cell membrane. TEM analyses of developing anther revealed that the elaioplasts in the mutant tapetum did not completely form till anther stage 12, suggesting that lipids might accumulate in the mutant tapetum (see results above). Other studies have reported that *CER3* is a core component of the complex required for synthesizing long-chain alkane and essential for the production of long-chain lipids (Bernard et al. 2012). Lipids biosynthesis might be affected in *cer3-8* mutant due to the mutation of *CER3* gene. Besides, excess tryphine was found on the surface of *cer3-8* pollen at maturity. So, disruption of *CER3* may influence biosynthesis of long-chain lipids, leading to *cer3-8* mutant phenotypes.

The reduction in 29-carbon and 30-carbon lipid level (Hannoufa et al. 1993; Preuss et al. 1993; the present study) is the common feature of all *cer* mutants. Hence, these long-chain lipid molecules are necessary for pollen-stigma interactions, either for directly signaling the stigma or for stabilizing other essential molecules. Further characterization of the *CER* gene products, including identification of their biochemical functions, should further elucidate the role of lipids in pollen-stigma communication, not only in *Arabidopsis*, but in other angiosperms as well.

## Acknowledgements

We thank the Salk Institute for providing seeds of *Arabidopsis* T-DNA insertion lines.

## Supporting Information

**Figure. S1.**
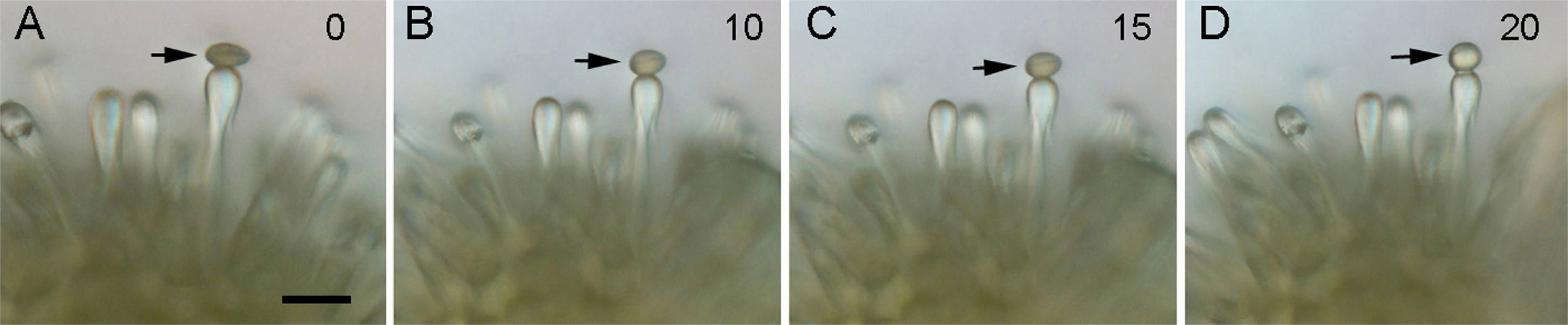
The hydration of *cer3* pollen on the *cer3* stigma was rescued under high humidity. Pollen expands (arrows) with shape changing from elliptic (A) to global (B-D) along with time course when placed on the *cer3* stigma surface, indicating that the *cer3* pollen is hydrated. Numbers (upper right corner) show times in minutes. The examples depicted here are representative result of more than 100 pollen grains. Bar = 100 μm in (A-D).

